# GeDi: Simplifying Gene Set Distances for Enhanced Omics Interpretation in R/Bioconductor

**DOI:** 10.1101/2025.10.10.681592

**Authors:** Annekathrin Silvia Nedwed, Arsenij Ustjanzew, Najla Abassi, Leon Dammer, Alicia Schulze, Sara Salome Helbich, Michael Delacher, Konstantin Strauch, Federico Marini

## Abstract

**Background:** Functional enrichment analysis is a standard component in many omics data analysis workflows, supported by a variety of methods and algorithms. However, despite their utility and wide application, these methods often return the results as an extensive and redundant list of gene sets, impeding interpretation and hypothesis generation. Moreover, network based information can provide additional biological context through functional interaction data, yet this is often overlooked by existing tools.

**Results:** We developed GeDi, an R/Bioconductor package designed to streamline and standardize the interpretation of functional enrichment results. GeDi aggregates gene sets into biologically meaningful clusters using a suite of gene set distance metrics and clustering algorithms, aimed to reduce redundancy and improve clarity. GeDi also enables the integration of protein-protein interaction (PPI) data, through the implementation of a weighted distance metric, providing a richer biological context by capturing functional connectivity between pathways and their components. The package offers visualizations, aggregation, and automated reporting, and is available as both a stand-alone R-package and an interactive Shiny application.

**Conclusion:** GeDi facilitates clearer, faster interpretation of enrichment results by combining clustering and network context. Application to a public RNA-seq dataset revealed coherent biological themes, supporting both experimental and computational research. GeDi is freely available in the Bioconductor project under the MIT license (https://bioconductor.org/packages/GeDi), and a demo instance is accessible on the Shiny server (http://shiny.imbei.uni-mainz.de:3838/GeDi).

## Background

One of the primary objectives in biological research is to uncover the molecular mechanisms driving phenotypes, traits and disease states (Cano-Gamez and Trynka 2020; Geistlinger et al. 2021). In bioinformatics, this is often done by means of functional enrichment analysis, which identifies pathways or biological processes that are statistically overrepresented in gene or protein sets defined by shared characteristics (e.g. differential expression or genetic variation) (Subramanian et al. 2005; Geistlinger et al. 2021; Garcia-Moreno, Adrian & López-Domínguez et al. 2022). As functional enrichment analysis became a fundamental step of modern omics workflows (Xie & Jauhari et al. 2021), various computational strategies have emerged, typically classified into over-representation analysis (ORA), functional class scoring (FCS), and pathway topology (PT)-based methods (Khatri, Sirota, and Butte 2012; Ma & Shojaie et al. 2019; Geistlinger et al. 2021). In general, ORA focuses on identifying gene sets with an overrepresentation of differentially expressed genes, while FCS methods examine the distribution of these genes within a ranked list (Subramanian et al. 2005). PT-based methods additionally incorporate pathway structure information to evaluate the impact of expression changes in genes and proteins (Ma & Shojaie et al. 2019).

In the broader scientific landscape, ORA methods remain the most widely adopted functional enrichment approaches due to their simplicity, efficiency, and broad applicability across biological research (Geistlinger et al. 2021). While pathway-topology methods offer deeper mechanistic insights by incorporating pathway structure (Khatri, Sirota, and Butte 2012), the widespread availability and support for ORA has established it as the standard in many workflows. Popular tools such as topGO (Alexa et al. 2006), g:Profiler (Kuleshov et al. 2016; Kolberg et al. 2023) and clusterProfiler (Wu et al. 2021) demonstrate this, and offer access to diverse annotation databases like Gene Ontology (GO), the Kyoto Encyclopedia of Genes and Genomes (KEGG), Reactome and the Molecular Signatures Database (MSigDB) (Ashburner et al. 2000; Liberzon et al. 2015; Fabregat et al. 2018; Kanehisa et al. 2023).

However, the output of most ORA methods typically consists of long, redundant gene lists, often complicating interpretation (Huang et al. 2009; Merico et al. 2010). Despite the existence of tools such as EnrichmentMap (Merico et al. 2010), GeneSetCluster (Ewing et al. 2020) or GScluster (Yoon et al. 2019), which focus on the aggregation of similar gene sets into clusters, interpretation is still mostly done manually due to the absence of biological context, which could be obtained from network data, such as protein-protein interactions (PPIs). While tools such as GScluster (Yoon et al. 2019) address this through the implementation of a PPI-weighted gene set distance score, the tool is limited by the amount of species supported in the analysis.

In this work, we present GeDi, an R/Bioconductor package designed to streamline the aggregation and interpretation of functional enrichment analysis results. This is achieved by computing **Ge**ne Set **Di**stance scores to quantify relationships between reported gene sets, which then serve as the basis for clustering to group similar sets, enhance clarity, and reduce redundancy. In order to accommodate a variety of different research scenarios, GeDi offers a range of distance metrics and clustering methods, as well as the integration of biological network data (e.g., PPI) to provide additional biological context. The package can be used programmatically or interactively via a Shiny app (provided within the package itself), using plain text files, spreadsheets or common R/Bioconductor data structures as input (Huber et al. 2015). This makes GeDi directly compatible with commonly used functional enrichment implementations, allowing for a seamless integration into various analysis workflows. As an essential downstream data exploration step, GeDi enables a more structured interpretation of results through visualization and aggregation of the large-scale enrichment results, ultimately reducing manual effort and facilitating hypothesis generation.

GeDi is freely available under the MIT license on Bioconductor (https://bioconductor.org/packages/GeDi), with the development version accessible on GitHub (https://github.com/AnnekathrinSilvia/GeDi) and a demo instance available at http://shiny.imbei.uni-mainz.de:3838/GeDi.

## Implementation

### General Design

The GeDi package is implemented in the R programming language within the Bioconductor project. We used the Shiny framework (Chang et al. 2024) to implement an application which can be run locally on the infrastructure of each individual user. Additionally, GeDi can be hosted on a server, as demonstrated in the demo instance (http://shiny.imbei.uni-mainz.de:3838/GeDi), making it suitable for internal group use by multiple users, or in situations where the exchange of sensitive personal data is necessary.

GeDi is designed to integrate seamlessly into existing omics workflows by facilitating the interpretation of functional enrichment results, such as those typically produced in bulk or single-cell RNA-seq analyses (Van den Berge et al. 2019; Amezquita et al. 2020; Ludt et al. 2022). To support a broad usability, GeDi accepts input in plain text, spreadsheets, R data.frames or as a GeneTonicList object (Marini et al. 2021) (Figure 1A). The only mandatory input requirement is a table with two columns named “Genesets” and “Genes”. While the format is flexible, we recommend using standardized identifiers such as HGNC (Seal et al. 2023), ENSEMBL (Yates et al. 2020), or Gencode (Frankish et al. 2019) for genes, and pathway IDs from GO (Ashburner et al. 2000), KEGG (Kanehisa et al. 2023), Reactome (Fabregat et al. 2018), or MSigDB (Liberzon et al. 2011) for gene sets.

**Figure 1.**
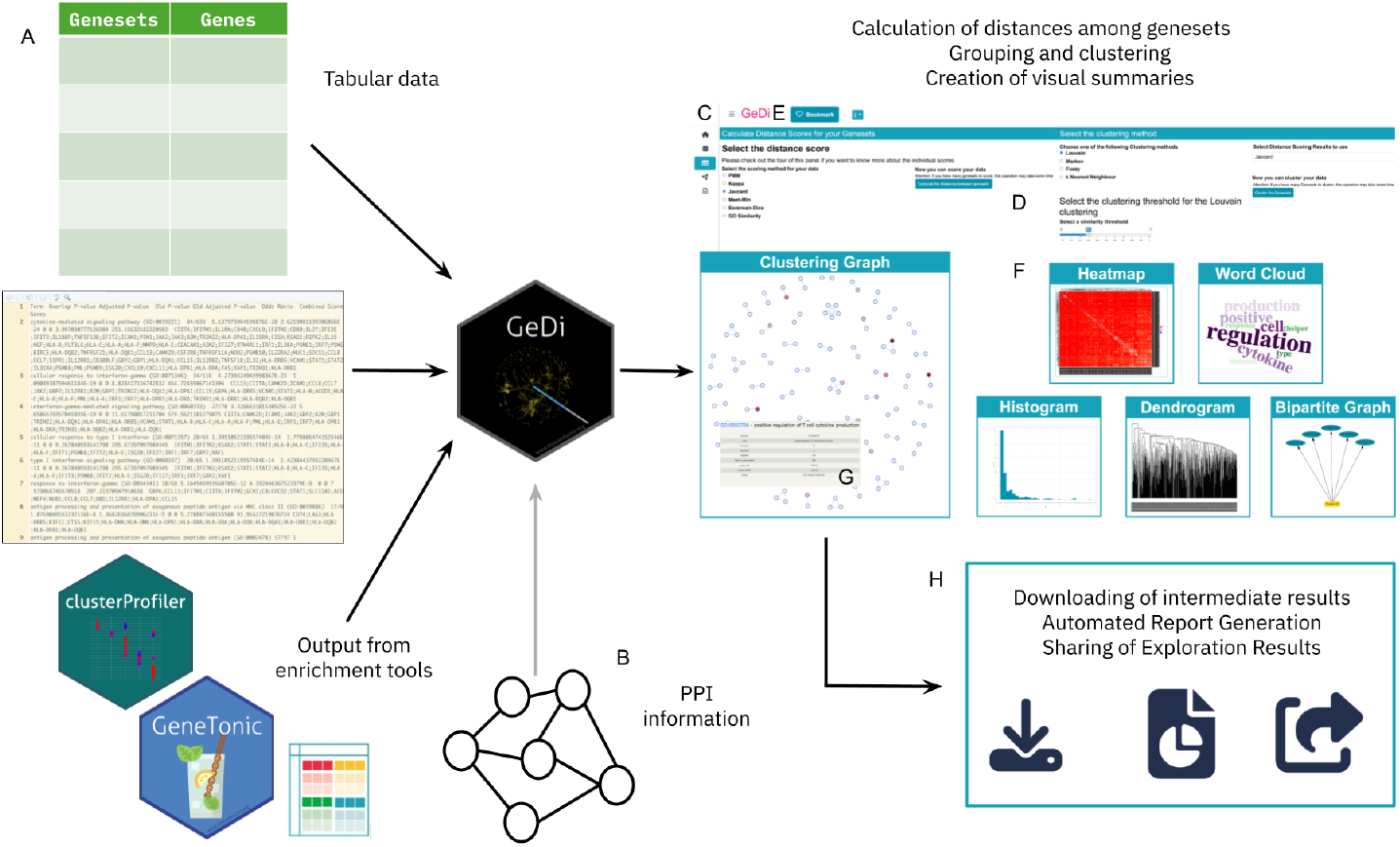
Overview of analysis workflow with GeDi. **A:** GeDi supports multiple input formats, including tabular data, enrichment tool outputs (e.g. topGO, clusterProfiler, etc.) and the GeneTonicList object format. **B:** Users may optionally provide their own PPI data instead of downloading it through the app. **C:** Overview of the individual interface panels. **D:** The user interface features intuitive selectors and buttons to guide users through the application. **E:** The Bookmark button allows marking of interesting genes and gene sets. **F:** GeDi offers a variety of visualisation options. **G:** A hover feature enables detailed inspection of gene sets. **H:** GeDi provides options to save intermediate results and generate a report of the exploration session for reproducibility and result sharing.

Additionally, GeDi can accept information on the protein-protein interactions as optional input (Figure 1B). While the PPI information can be fetched and cached locally during runtime, providing the data upon launching of the app can reduce manual intervention, thus enhancing the reproducibility of the analysis.

The GeDi package provides nearly all of its functionality through a single entry point: the GeDi() function. This launches a web application that guides users through the exploration of functional enrichment results using a clear, section-based layout (Figure 1C). Each panel of the app focuses on a specific aspect of the data, offering tailored visualizations and summaries. The interface is built using the bs4Dash (Granjon 2024) package, ensuring a modern, responsive user experience.

To support intuitive onboarding and enhance user engagement, GeDi includes interactive, guided tours implemented with the rintrojs package (Ganz 2016). These tours accompany users through the different components of the app, providing context-sensitive help and highlighting individual UI elements through modals.

To prevent overwhelming users, the app design emphasizes clarity and flow, with panels using collapsible elements, often revealed sequentially. This structure not only makes the app easier to use but also supports a more effective, step-by-step exploration of complex datasets (Figure 1D).

Users can provide input data at the beginning of a session, or interactively during usage. The application also features a bookmarking system, allowing users to flag genes and gene sets of interest throughout their exploration (Figure 1E). Bookmarked items are summarized in a dedicated Report panel and automatically included in the downloadable report generated at the end of the session. Buttons for bookmarking are consistently located in the panel headers, making the process straightforward.

While designed for interactive use, most functions are also exported and can be reused in scripted analyses (e.g., within RMarkdown or Jupyter notebooks), making GeDi flexible for both hands-on exploration and reproducible workflows.

### Typical usage workflow

GeDi requires as input the results of a functional enrichment analysis, comprising at least two columns named “Genesets” and “Genes”. These can be supplied directly via the function call GeDi() with a corresponding data.frame, or uploaded through the *Data Upload* panel in the GeDi interface, supporting plain text, spreadsheet, or RDS formats.

GeDi can be readily integrated as a downstream step in workflows that include functional enrichment analysis, such as for example (bulk) RNA-sequencing data analysis workflows such as those demonstrated in Additional File 1 and 2 of the Supplementary Material.

In addition to the core input, GeDi optionally accepts a matrix of protein-protein interaction scores (ppi_df) and a distance matrix between gene sets (distance_scores). Both can be generated within GeDi but may incur significant computation time; saving and reusing these objects is therefore advisable. Download options for these are available within the interface. To facilitate broad applicability, GeDi currently supports outputs from widely used R packages for enrichment analysis, including topGO (Alexa et al. 2006) and clusterProfiler (Wu et al. 2021). Alternatively, users are encouraged to follow the GeneTonic workflow (Marini et al. 2021), which would generate and preprocess results in a format that is directly supported by GeDi.

While GeDi only requires gene set and gene information, the inclusion of additional metadata from enrichment analysis tools — such as gene set descriptions, p-values, or set sizes — can significantly enhance the user experience, e.g., through coloring of the visualisations according to the metadata (Figure 1F-G). For example, such metadata enables richer visualizations, including node coloring in graphs, and improves interpretability within the *Clustering Graph* panel. GeDi supports iterative data exploration through a range of interactive visual summaries, including heatmaps, gene set graphs, bipartite graphs, and word clouds (Figure 1F-G). The computed distance scores serve as the basis for clustering, helping to group related gene sets and reveal broader functional patterns. Users can also hover on the visualisations in GeDi to explore additional information, for example through dynamic links to external databases such as AmiGO (The Gene Ontology Consortium 2019), KEGG (Kanehisa et al. 2023), or Reactome (Fabregat et al. 2018), which support deeper biological interpretation.

Although GeDi is primarily accessed via its interactive web interface, all functions operate on standard R objects, allowing straightforward integration into existing R/Bioconductor workflows (e.g., (Huber et al. 2015; Van den Berge et al. 2019; Ludt et al. 2022). A recommended final step in a typical analysis with GeDi is the generation of an HTML report, which facilitates both result sharing and computational reproducibility (Figure 1H).

## Results

### GeDi implements distances and clustering approaches to summarize enrichment results

GeDi facilitates the interpretation of functional enrichment results by grouping gene sets into functionally coherent clusters based on their pairwise (dis)similarity. This is achieved through the computation of gene set distance scores, which quantify how similar or dissimilar gene sets are in terms of gene content, semantic meaning, or network context. GeDi implements a range of distance metrics, which can be broadly categorized into: (1) **set-based** metrics (Jaccard (Jaccard 1912; Levandowsky and Winter 1971), Sørensen-Dice (Sorensen 1948), Kappa (Cohen 1960), and Meet-Min (MM) (Yoon et al. 2019)), relying solely on gene set overlap; (2) **hybrid** metrics (protein-protein interaction weighted Meet-Min distance score (pMM, (Yoon et al. 2019))), which combine gene set overlap with protein–protein interaction (PPI) context; and (3) **semantic** metrics (GO dissimilarity (Yu et al. 2010)), which exploit the structure of the Gene Ontology to assess functional similarity. Each metric captures a different aspect of relatedness: while set-based scores are sensitive to direct gene set overlap, pMM adds biological relevance by considering interaction networks, and semantic metrics provide a broader, ontology-driven similarity measure. Based on the computed distances, GeDi applies clustering algorithms to aggregate gene sets into groups that represent shared biological functions. Available algorithms include Louvain (Blondel et al. 2008) and Markov (van Dongen 2000; Enright, Van Dongen, and Ouzounis 2002) clustering, which detect communities in similarity networks; PAM (Partitioning Around Medoids (Kaufman and Rousseeuw 1987)), a centroid-based method; and Fuzzy clustering (Huang et al. 2007; Huang et al. 2009), which allows overlapping cluster membership. The choice of clustering algorithm affects the structure and granularity of the resulting clusters: Louvain and Markov typically yield discrete, non-overlapping groups, while Fuzzy clustering can capture functional cross-talk by assigning gene sets to multiple clusters. These complementary options enable users to explore different aspects of the functional landscape represented in enrichment results. A detailed description and definition of the distance scores as well as of the clustering algorithms can be found in Additional File 3.

### Comparison of Meet-Min score and pMM score

In a further inspection and analysis of the methods and functionality provided by GeDi, we inspected how the additional network information incorporated in the pMM can leverage the score and results. For this purpose, we investigated how a random removal of genes from the gene sets would influence the resulting scores and clusters.

Hence, we first removed one to 15 genes from gene sets with more than 10 genes, removing at most half of the genes in any given set. Then we scored the data using either the pMM or the MM score. Afterwards, we iterated 100 times the community-detection Louvain clustering on the original data as well as on the randomly reduced data, as this algorithm is inherently non-deterministic. Afterwards, we compared the clustering results using the Adjusted Rand Index (ARI) coefficient as a measure of similarity between two data clusterings (Rand 2012). In this specific use case, we used the ARI coefficient to compare the clustering results of the reduced datasets to the original clustering result on the whole set of gene sets. To compare the two distance metrics, we plotted the resulting ARI scores for the MM and pMM score, which can be observed in Supplementary Figure 1. Upon closer inspection, we observed that the pMM ARI is almost exclusively higher than the MM ARI score. This indicates that the network information included in the pMM score can help in retaining the information that would be lost by the removal of genes, which would significantly impact the computed MM scores.

### GeDi enables the quantification and navigation of gene set similarities after functional enrichment analyses

Following the differential expression and functional enrichment analysis of various omics workflows, GeDi provides an ideal additional step in facilitating the exploration and interpretation of the resulting enriched pathway data. The functional enrichment results of several tools can be directly provided as input to GeDi, provided that they follow the expected input format of the package. However, if the input format does not adhere to the requirements, GeDi also implements functionalities to transform and adapt it.

As described above, GeDi provides a variety of gene set distance scores to quantify their (dis)similarity between individual gene sets. In addition, to represent these calculated distance scores, GeDi offers three main visualisation types: a heatmap, a dendrogram, and a network-like representation. The heatmap offers a complete view of all pairwise distances, allowing users to detect global patterns through color gradients, while the dendrogram adds a hierarchical perspective revealing nesting groups. Lastly, in the network-like representation, gene sets are represented as nodes and edges are drawn between gene sets with a distance below a user defined threshold. The network can highlight functional modules and local neighbourhoods, whose granularity can be adjusted through the threshold parameter. Together, these visualisation options provide some initial global and focused interpretation of the overarching patterns of regulation in the enrichment results.

These patterns can be further inspected through the application of one of the available clustering algorithms in GeDi. The results of the clustering process can be visualised in different ways such as in a graph network, a bipartite graph, or word clouds. For the graph network, nodes represent gene sets and edges are drawn between gene sets belonging to the same cluster, while in the bipartite graph, nodes represent gene sets and clusters, with edges connecting gene sets to their respective cluster(s). Lastly, in the word cloud representation, users can select a specific cluster and the word cloud will visualise the most prominent biological terms associated with the gene sets in the cluster. However, this is dependent on the availability of a (biological) description of each gene set in the input meta data.

Additionally, GeDi provides functionality to export the exploration session in a ready-to-share HTML report. This report includes tables of the original input data, as well as the selected parameters during the session (such as distance scoring method, clustering algorithm and thresholds) as well as the visualisation of the calculated distance scores and clustering results. This report does not only serve as a documentation on the conducted exploration, but also as a means to share and reproduce the analysis.

### Applying GeDi to publicly available data to enhance the interpretation of functional enrichment results

We use a publicly available bulk RNA-seq dataset, published by Delacher et al. (Delacher et al. 2020) (GEO accession: GSE130842; https://www.ncbi.nlm.nih.gov/geo/query/acc.cgi?acc=GSE130842), to demonstrate GeDi’s functionality. The dataset comprises 56 samples from various murine tissues, including distinct regulatory T cell (Treg) subpopulations. We focused on Klrg1^−^Nfil3(GFP)^−^ and Klrg1^+^Nfil3(GFP)^+^ Tregs, yielding a subset of eight samples. For the preprocessing, we followed the workflow proposed by (Ludt et al. 2022), documented in Additional File 1 and on Github (https://github.com/AnnekathrinSilvia/GeDi_supplement). Ultimately, we generated a GeneTonicList object (Marini et al. 2021), which we used as an input for GeDi.

### Providing the data to GeDi

We initiated the analysis by launching GeDi via the GeDi() command and loading the prepared GeneTonicList object into the launched app. In a preprocessing step, we first filtered out GO terms associated with ≥ 200 genes, reducing the total from 500 to 421 gene sets. Subsequently, we retrieved the protein-protein interaction matrix from STRING (version 12.0.0), using GeDi’s built-in functions. A detailed workflow of this can be found in Additional File 1 or on Github (https://github.com/AnnekathrinSilvia/GeDi_supplement).

### Quantifying gene set similarity using gene set distance scores

In this study, we systematically evaluated all distance metrics available in GeDi, ultimately focusing on the pMM and GO semantic distances, which consistently produced medium-sized, biologically meaningful clusters. In Figure 2, we visualised the resulting distance scores using the heatmap visualisation available in GeDi. Both heatmaps show that independent of the used distance metric, most of the gene sets had a high distance close to 1, with distinct clusters of smaller distances visible along the diagonal (Figure 2A-B). However, the pMM heatmap (Figure 2A) revealed more concentrated blocks of similar gene sets, especially in the upper-left and lower-right corners of the heatmap, compared to the GO heatmap (Figure 2B). This was already a first indicator of the additional information gained through the incorporation of network-information from PPI-networks.

**Figure 2.**
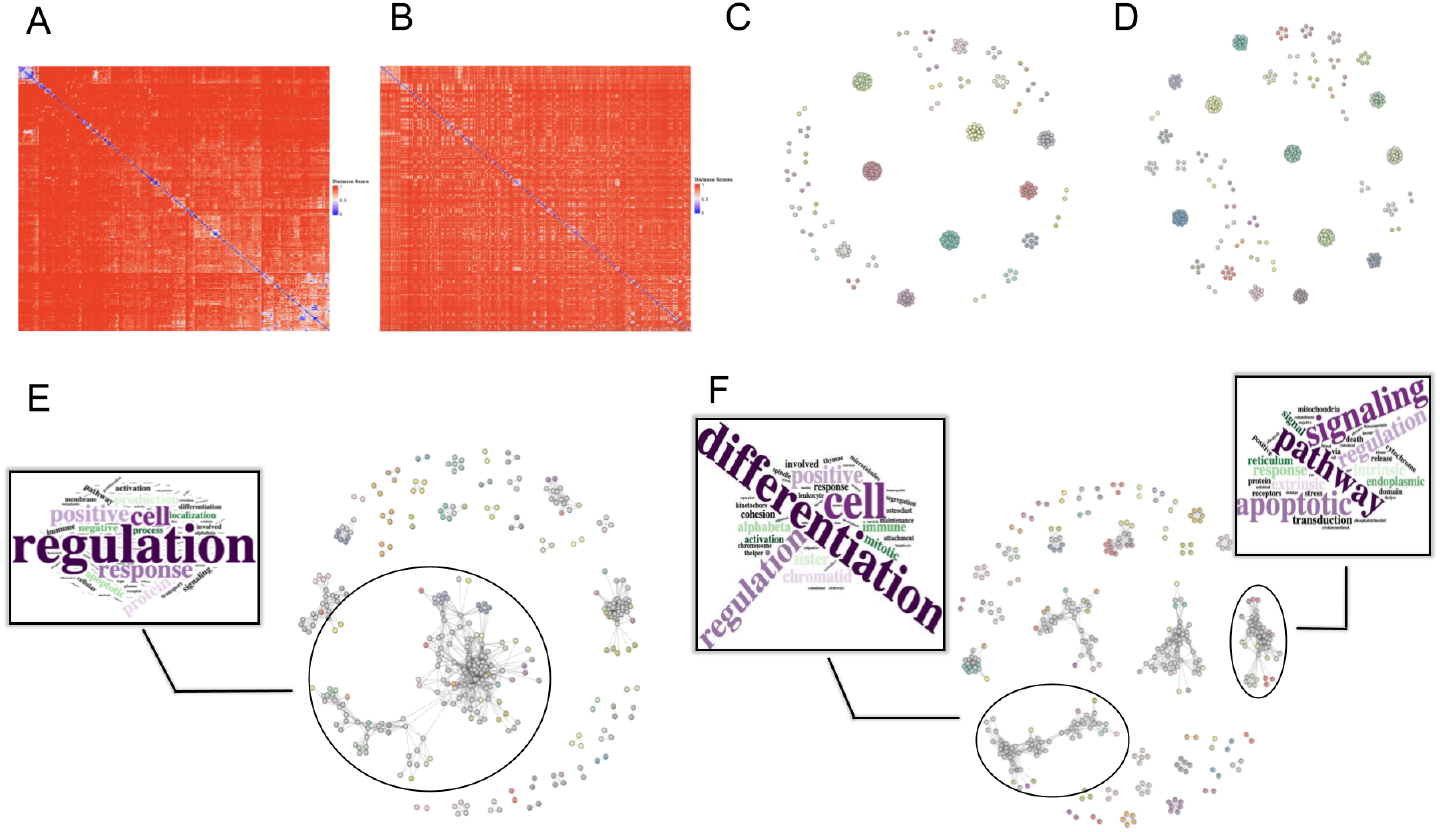
Exemplary output generated with GeDi based on pMM and GO distance score for the Delacher et al. dataset. **A:** Heatmap of the calculated pMM distance scores. **B:** Heatmap of the calculated GO distance scores. **C:** Louvain clustering results based on the pMM distance scores. **D:** Louvain clustering results based on the GO distance scores. **E:** Fuzzy clustering results based on the pMM distance scores. **F:** Fuzzy clustering results based on the GO distance scores.

### Aggregating gene sets into similar groups using clustering algorithms

Following the distance computation, we selected two clustering algorithms, Louvain and Fuzzy clustering, to showcase the capabilities of GeDi. For the thresholds, we consistently chose 0.5 for the Louvain clustering as well as the Fuzzy similarity, membership and clustering threshold.

In Figure 2, the resulting clustering graphs of the two clustering approaches can be seen, with the clustering of the pMM scores in Figure 2C and Figure 2E and the GO distances in Figure 2D and Figure 2F. We can observe that the Louvain clustering algorithm resulted in similar numbers of clusters for both the pMM and GO scores (41 vs. 49 clusters, Figure 2C-D). Both graphs show an overall composition of smaller clusters, with some larger clusters in between. Using the hovering tooltip of GeDi (Figure 1G), we identified that these larger clusters captured biological coherent themes, including T cell differentiation, cell division, and protein localization.

The Fuzzy clustering on the other hand resulted in an overall higher number of clusters (127 clusters for both distance metrics), of which many shared cluster members. The largest clusters in the middle of the graphs showed to be enriched with genes involved in immune regulation and T cell response (Figure 2E). We could also find biological processes of T cell differentiation and apoptotic signaling pathways in these clusters, which was more prominently observable in the clustering results on the GO distances, where these biological processes were split into two larger clusters (Figure 2F).

Overall, the recurring biological themes - most prominently, T cell differentiation and proliferation, as well as protein localisation and apoptotic signaling pathways - align well with the original findings and biological context of the used dataset. Delacher et al. (2020) identified Klrg1^−^Nfil3^−^ and Klrg1^+^Nfil3^+^ Treg cells as developmental stages within the lineage of tissue-resident Tregs, a subpopulation involved in tissue regeneration (Ito et al. 2019; Delacher et al. 2020). The observed enrichment of differentiation- and proliferation-related pathways reflects the transition from Klrg1^−^Nfil3^−^ to Klrg1^+^Nfil3^+^ late precursors cells, driven by increased expression of Klrg1 and Nfil3. Concurrently, changes in surface protein and antigen expression during this process explain the strong representation of pathways linked to protein localization and transport. With the help of GeDi, we were able to enhance and highlight these biological processes inherent in the data, ultimately allowing for a facilitated and refined interpretation of the original functional enrichment data.

## Discussion

Functional enrichment remains a fundamental step of modern omics analysis pipelines, yet the interpretability of its results continue to pose challenges. Standard methods often yield long, redundant gene set lists that obscure biological signals, ultimately hindering hypothesis generation. While similarity- and clustering-based summarization approaches exist, they are frequently limited in scope or interactivity (Supplementary Table 1), and many enrichment tools still rely on manual inspection, lacking meaningful integration of biological context. Additionally, in a recent article, Wijesooriya et al. (Wijesooriya et al. 2022) found that researchers tend to use these tools in an improper manner, while also not reporting methodological details appropriately.

GeDi addresses these limitations by offering a unified framework that integrates multiple gene set distance metrics, various clustering algorithms, and interactive visualisation tools to help users explore and interpret enrichment results in a structured manner. While we currently support a broad range of metrics, the modular architecture of GeDi will allow us to incorporate future methods, ensuring long-term compatibility and relevance as enrichment methodologies evolve.

Designed with both flexibility and usability in mind, GeDi can be used programmatically within notebooks and scripts, or interactively through its Shiny application. The app offers powerful drilldown capabilities that allow users to interrogate specific gene sets, explore annotation details, and access external resources, facilitating deeper biological interpretation.

Sessions can be concluded with automated report generation, enabling reproducible documentation of each analysis step. Intermediate results are passed along transparently, allowing users to resume or branch off from any point in the workflow. This design supports both exploratory and structured research scenarios, which has been shown essential in other works (Marini et al. 2021; Ludt et al. 2022) and can be seen as a democratizing factor in interdisciplinary efforts.

While there is a wide variety of exploration tools for enrichment results, such as for example EnrichmentMap (Merico et al. 2010), most of these tools only offers functionality to visualise the results, without any functionality offered to reduce the complexity of the result list (Supplementary Table 1). While there are other tools, such as GScluster or GeneSetCluster, which have addressed these aspects of gene set aggregation (Supplementary Table 1), they often impose stricter input requirements or lack integration with the broader R/Bioconductor ecosystem. In contrast, GeDi requires minimal input - only a two-column table of genesets and genes - yet seamlessly accepts widely used outputs (e.g. from topGO, clusterProfiler or other apps such as GeneTonic which aim to reuse standardized objects and classes). This positions GeDi as an accessible and robust post-enrichment summarization tool for a broad user base.

Looking forward, we aim to extend GeDi’s applicability to GSEA-style enrichment results, further broadening its scope. Expanding supported sources of PPI and functional interaction data is another key direction, with the goal of offering richer network-based summaries.

Given the increasingly widespread adoption of tools based on large language models and single cell foundation models for the interpretation of biological data, we anticipate that GeDi can provide better suited summaries that would in turn enhance the answers returned by such agents (Cui et al. 2024; Li et al. 2025).

In summary, GeDi serves as a scalable, user-friendly solution for the systematic summarization of enrichment results, efficiently bridging the gap between statistical detection and meaningful biological insight.

## Conclusion

In our work we presented GeDi, a new R package developed to streamline and simplify the analysis and interpretation of functional enrichment results. GeDi provides several different distance metrics, which can be used to quantify the similarity of the gene sets in the input functional enrichment results, as well as a variety of clustering algorithms to aggregate the gene sets to groups based on their distance scores.

GeDi can reduce the extensive lists of gene sets arising from enrichment analyses to compact sets of functional clusters, fostering via different distance measurements and clustering approaches the careful inspection of various aspects of the data at hand. We expect that this approach, well rooted in the Bioconductor ecosystem of packages yet simple to adopt for many omics analysis workflows, will empower many scientists to generate viewpoints for clearer and better hypothesis generation and interpretation.

## Availability and Requirements

Project name: GeDi.

Project home page: https://bioconductor.org/packages/GeDi (release),

https://github.com/AnnekathrinSilvia/GeDi (development version).

Archived version: 10.5281/zenodo.16315835, package source as gzipped tar archive of the version reported in this article.

Project documentation: rendered at https://annekathrinsilvia.github.io/GeDi/.

Operating systems: Linux, Mac OS, Windows. Programming language: R.

Other requirements: R-4.4.0 or higher, Bioconductor 3.19 or higher. License: MIT.

Any restrictions to use by non-academics: none.

## Abbreviations

ARI: Adjusted Rand Index
FCS: Functional Class Scoring
GEO: Gene Expression Omnibus
GO: Gene Ontology
GSEA: Gene Set enrichment analysis
HGNC: HUGO (Human Genome Organisation) Gene Nomenclature Committee
KEGG: Kyoto Encyclopedia of Genes and Genomes
MSigDb: Molecular Signatures Database
ORA: Overrepresentation Analysis
PPI: Protein-Protein Interaction
RNA-seq: RNA sequencing
Treg: regulatory T cells

## Declarations

### Ethics approval and consent to participate

Not applicable

### Consent for publication

Not applicable

### Availability of data and materials

The datasets used in this manuscript and its supplements are available from the following articles: The dataset on the regulatory T cell subpopulations is included in PubMed ID: 31924477 (https://doi.org/10.1016/j.immuni.2019.12.002). Dataset deposited at the Gene Expression Omnibus (GEO) (GSE130884, project id: PRJNA541901) and accessed from the https://github.com/AnnekathrinSilvia/GeDi_supplement repository. Clinical trial number: not applicable.

The GeDi package can be downloaded from its Bioconductor page https://github.com/AnnekathrinSilvia/GeDi_supplement/ contains the code used to generate the supplemental material and the required input data to replicate the analyses presented in the use cases.

## Authors’ contributions

ASN - conceptualization, data curation, formal analysis, methodology, project administration, resources, software, visualization, writing - original draft, writing - review and editing. AU - data curation, software, writing - review and editing. NA - data curation, software, writing - review and editing. LD - software, writing - review and editing. AS - data curation, software, writing - review and editing. SSH - data curation, interpretation, writing - review and editing. MD - data curation, interpretation, writing - review and editing. KS - conceptualization, funding acquisition, resources, supervision, writing - original draft, writing - review and editing. FM - conceptualization, data curation, formal analysis, funding acquisition, methodology, project administration, resources, software, supervision, visualization, writing - original draft, writing - review and editing. All authors read and approved the final version of the manuscript.

## Acknowledgements

This work has been supported by the computing infrastructure provided by the Core Facility Bioinformatics at the University Medical Center Mainz, used also for deploying the demo instance. The authors thank the Bioconductor community and the attendees of the EuroBioC2024 for valuable feedback and suggestions.

## Authors’ information

### Authors and Affiliations

Institute of Medical Biostatistics, Epidemiology and Informatics (IMBEI), University Medical Center of the Johannes Gutenberg University Mainz, Rhabanusstr. 3, 55118, Mainz, Germany Annekathrin Silvia Nedwed, Arsenij Ustjanzew, Najla Abassi, Leon Dammer, Alicia Schulze, Konstantin Strauch & Federico Marini

Center for Thrombosis and Hemostasis (CTH), University Medical Center of the Johannes Gutenberg University Mainz, Langenbeckstr. 1, 55131, Mainz, Germany Federico Marini

Institute of Immunology, University Medical Center of the Johannes Gutenberg University Mainz, Langenbeckstr. 1, 55131, Mainz, Germany

Sara Salome Helbich & Michael Delacher

Research Center for Immunotherapy (FZI), University Medical Center of the Johannes Gutenberg University Mainz, Langenbeckstr. 1, 55131, Mainz, Germany

Sara Salome Helbich, Michael Delacher & Federico Marini

## Corresponding author

Correspondence should be addressed to Annekathrin Silvia Nedwed and Federico Marini.

