## Supplementary Figure 1 for "GeDi: Simplifying Gene Set Distances for Enhanced Omics Interpretation in R/Bioconductor"

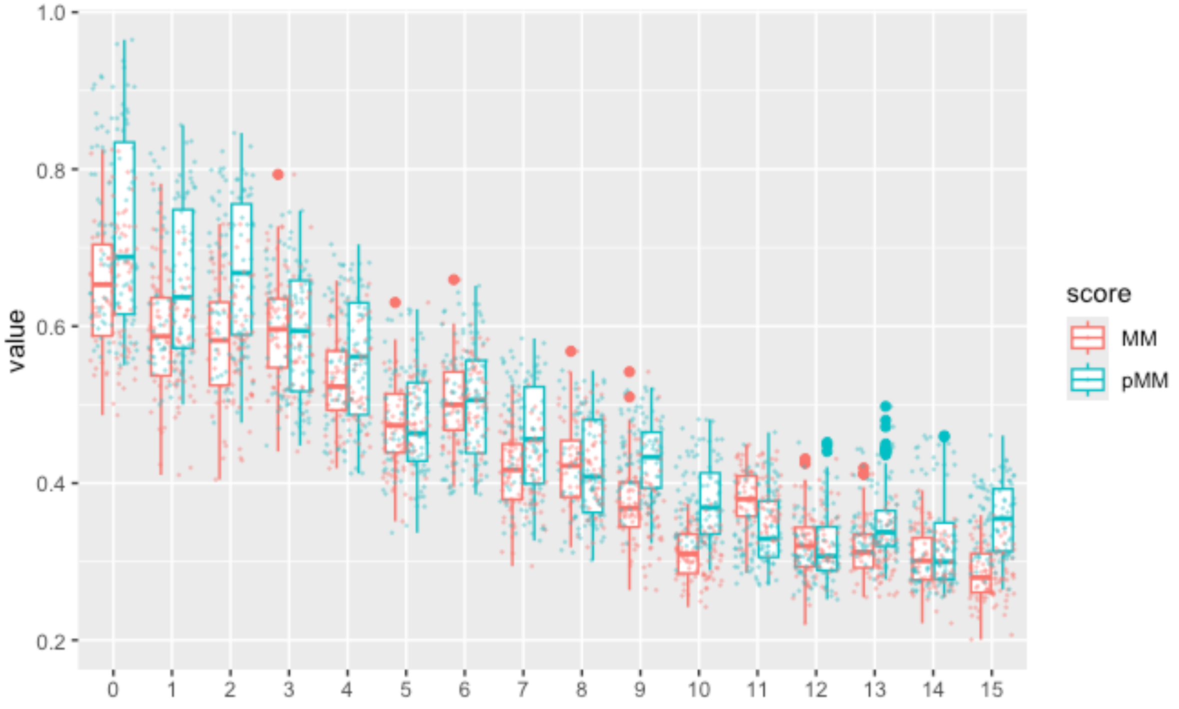


Supplementary Figure 1: The figure shows the box plot comparison of the ARI scores of the Louvain clustering results based on the MM and pMM distance scores. The x-axis shows the number of genes removed from the gene sets, while the y-axis shows the value of the ARI score.
