## Supplementary Materials and Methods for "GeDi: Simplifying Gene Set Distances for Enhanced Omics Interpretation in R/Bioconductor"

### Details on the Distance Scores implemented in GeDi

#### Meet-Min (MM) distance

In various biological research areas, the Meet-Min (MM) distance score is used to quantify the functional similarity between two sets such as gene sets or pathways (Yoon et al. 2019). By considering both the overlap and the relative sizes of the sets, the metric provides a quantitative measure of similarity. The Meet-Min distance score is then defined as

$$\mathrm{MM}(A,B) = 1 - \frac{|A \cap B|}{min(|A|,|B|)}$$

where $A$ and $B$ represent two distinct gene sets, respectively, and $|\cdot|$ represents their cardinality (i.e., the number of elements in the gene set).

The MM score is symmetric (i.e., $\mathrm{MM}(A,B) = \mathrm{MM}(B,A)$), with the range of the resulting distances being in $[0,1]$. A distance of $0$ denotes completely identical sets, whereas a distance of $1$ reflects completely disjoint sets (Nedwed 2025).

#### Cohen’s Kappa distance

The Cohen's Kappa distance score, hereafter referred to as Kappa distance score, is derived from the equally named Cohen's Kappa coefficient. Initially introduced to measure inter-rater reliability for categorical items (Cohen 1960), it has since been widely used as a quantitative measure of (dis)similarity between gene sets, e.g., in the work by (Yoon et al. 2019) or the Database for Annotation, Visualisation, and Integrated Discovery (DAVID) (Huang, Sherman, and Lempicki 2009; Huang et al. 2007).

As a distance measure, the Kappa distance quantifies the level of agreement between two sets and is defined as

$$\mathrm{Kappa}(A,B) = 1 - \frac{O - E}{1 - E}$$

with $O = \frac{|A \cap B| + |(A \cup B)^c|}{|U|}$

and $E = \frac{|A||B| + |A^c||B^c|}{|U|^2}$

In this score, $U$ is the union set of all genes (Yoon et al. 2019).

The Kappa distance score is symmetric (i.e., $\text{Kappa}(A, B) = \text{Kappa}(B, A)$) and the range of the resulting distances is $[0, 2]$. A value of $0$ indicates that the sets are completely identical, whereas a score of $2$ reflects completely disjoint sets.

Since other distance metrics used in this work result in distances in the interval $[0, 1]$, the Kappa distance score is transformed to match this range, ensuring comparability across all metrics. To normalise the Kappa distance score to the $[0, 1]$ range, the following formula was applied:

$$\text{Normalised\_Kappa}(A, B) = \frac{\text{Kappa}(A, B) - min}{max - min}$$

where $min$ and $max$ represent the smallest and largest possible Kappa distances, respectively. Given that these values are defined as $0$ and 1, the normalization simplifies to

$$\text{Normalised\_Kappa}(A, B) = \frac{\text{Kappa}(A, B)}{2}$$

This ensures all Kappa distances are scaled to the $[0, 1]$ interval (Nedwed 2025).

#### Jaccard distance

The Jaccard distance score is a measure of dissimilarity between two sets based on the commonly used Jaccard index (Jaccard 1912; Levandowsky and Winter 1971) . The Jaccard index measures the similarity between two sets by dividing the size of their intersection by the size of their union and is defined as

$$\text{Jaccard}(A, B) = 1 - \frac{|A \cap B|}{|A \cup B|}$$

The Jaccard distance score is symmetric (i.e., $\text{Jaccard}(A, B) = \text{Jaccard}(B, A)$), and the range of the resulting distances is $[0, 1]$. A distance of $0$ indicates that the sets are completely identical, while a distance of $1$ reflects completely disjoint sets.

#### The Protein-Protein Interaction weighted Meet-Min (pMM) distance

The Protein-Protein Interaction weighted Meet-Min (pMM) distance score is derived from the previously discussed Meet-Min distance score. Introduced in the work by authors (Yoon et al. 2019), the pMM score incorporates protein-protein interaction data to enhance the MM score by weighing it with functional information at the protein level. While the MM score measures the overlap between gene sets, the pMM score adds a PPI-based factor, recognising that gene overlap alone may not fully capture functional relationships between biological processes. Since functional interactions often occur at the protein level, the additional PPI component provides a more nuanced measure of similarity between gene sets.

The pMM distance score is defined as

$$\text{pMM}(A, B) = min(\text{pMMlocal}(A \to B), \text{pMMlocal}(B \to A))$$

With

$$\text{pMMlocal}(A \to B) = 1 - \frac{|A \cap B|}{min(|A|, |B|)} - \frac{\alpha}{min(|A|, |B|)} \sum_{a \in A-B} \frac{w \sum_{b \in A \cap B} P(a, b) + \sum_{b \in B-A} P(a, b)}{max(P)(w|A \cup B| + |B - A|)}$$

$P$ is a PPI matrix, a numerical matrix indicating a possible interaction of two proteins through confidence scores in the range of $[0, 1]$. $P(a, b)$ is the interaction score of two proteins, represented by their coding genes $a$ and $b$. $\alpha$ is a scaling factor to control the influence of the protein interactions on the resulting distances, with the range $[0, 1]$. If set to $0$, the pMM score is identical to the MM score, while a value of $1$ balances the influence of the two components of the score. Lastly

$$w = \begin{cases} \frac{|A|}{|A|+|B|}, \text{ if } |A| \leq |B| \\ \frac{|B|}{|A|+|B|}, \text{ otherwise} \end{cases}$$

and $\text{pMMlocal}(B \to A)$ is symmetrically defined.

The resulting pMM distance score of two gene sets $A$ and $B$ is symmetric (i.e., $\text{pMM}(A, B) = \text{pMM}(B, A)$) with a range of $[0, 1]$. A value of $0$ indicates that the sets are either completely identical or have a high degree of functional similarity based on the provided interaction information, whereas a distance of $1$ indicates completely disjoint sets (Nedwed 2025).

#### Sørensen-Dice distance

The Sørensen-Dice distance (Sorensen 1948) is a metric based on the corresponding similarity metric which is used to quantify the similarity between two sets. It is defined as

$$Sorensen - Dice(A, B) = 1 - \frac{2 * |A \cap B|}{|A| + |B|}$$

The Sørensen-Dice distance is particularly useful when dealing with binary or categorical data, making it versatile across various domains where set-based or presence/absence comparisons are essential.

#### GO Semantic Similarity

The GO semantic similarity score is based on the implementation of various measures for semantic similarities implemented in the GOSemSim R package (Yu et al. 2010). The package implements four different information content and as well as two graph-based similarity measures.

In the GeDi app, the Wang method that is based on the graph topology of the GO database is used as the default method (Yu et al. 2010). However, when using the function stand-alone in a scripted analysis, all different similarity measures can be used.

For a more detailed explanation of the various measures available, we refer interested readers to either (Nedwed 2025) or (Yu et al. 2010).

### Details on the clustering algorithms implemented in GeDi

GeDi provides multiple clustering algorithms to group functionally related gene sets based on their similarity scores. Each algorithm has distinct characteristics suited for different analysis needs.

The Louvain algorithm is a modularity-based community detection method that identifies densely connected groups of gene sets within a network. It follows a hierarchical two-phase process, first merging nodes into local communities and then optimizing modularity recursively on the condensed network. Due to its efficiency and scalability, Louvain clustering is well-suited for large biological datasets (Blondel et al. 2008). In contrast, the Markov Clustering (MCL) algorithm simulates random walks in a graph to identify clusters. Through alternating steps of expansion, which spreads flow across connected nodes, and inflation, which strengthens high-probability paths while suppressing weaker ones, MCL effectively detects modular structures in biological networks (van Dongen 2000; Enright, Van Dongen, and Ouzounis 2002).

For cases where gene sets may belong to multiple functional groups, Fuzzy Clustering, as implemented in the DAVID database, allows nodes to be assigned to multiple clusters based on Kappa distance scores. This method reflects the reality that genes often participate in multiple biological processes, providing a more nuanced interpretation of functional enrichment results. Initial clusters are formed based on functional similarity, and iterative merging refines the final clusters to ensure coherence (Huang et al. 2007; Huang, Sherman, and Lempicki 2009). Additionally, Partitioning Around Medoids (PAM) offers a robust alternative by selecting representative medoids within each cluster to minimize intra-cluster dissimilarity. Unlike K-means, PAM is more resistant to noise and outliers, making it particularly useful for gene set clustering where variability is high (Kaufman and Rousseeuw 1987).

By integrating these diverse clustering methods, GeDi provides a flexible framework for organizing enrichment results, allowing researchers to select the most appropriate approach based on dataset characteristics and analytical requirements. For a more detailed discussion of these clustering algorithms and their implementation within GeDi, interested readers are referred to (Nedwed 2025).
