## Additional File 1 and 2 for "GeDi: Simplifying Gene Set Distances for Enhanced Omics Interpretation in R/Bioconductor": GeDi_Additional_File_1.html

Using GeDi on the T cell dataset (GSE130842)


Code 

- Show All Code
- Hide All Code
- Download Rmd

### Using GeDi on the T cell dataset (GSE130842)

###### Annekathrin Silvia Nedwed

Institute of Medical Biostatistics, Epidemiology and Informatics (IMBEI), Mainz   
  
###### Arsenij Ustjanzew

Institute of Medical Biostatistics, Epidemiology and Informatics (IMBEI), Mainz   
  

###### Sara Salome Helbich

Institute of Immunology, University Medical Center Mainz, Mainz, Germany   
Research Center for Immunotherapy (FZI), Mainz, Germany   
  

###### Michael Delacher

Institute of Immunology, University Medical Center Mainz, Mainz, Germany   
Research Center for Immunotherapy (FZI), Mainz, Germany   
  

###### Konstantin Strauch

Institute of Medical Biostatistics, Epidemiology and Informatics (IMBEI), Mainz   
  

###### Federico Marini

Institute of Medical Biostatistics, Epidemiology and Informatics (IMBEI), Mainz   
Research Center for Immunotherapy (FZI), Mainz, Germany   
Center for Thrombosis and Hemostasis (CTH), Mainz   
  
###### 16 July 2025

### 1 About the data

The data illustrated in this document is an RNA-seq dataset, available at the Gene Expression Omnibus under the accession code GSE130842.

The data represents a mouse data set of 32 different samples across different tissues and conditions. The data is part of a manuscript to analyse different tissue regulatory T cell populations (Delacher et al. 2020) - the manuscript is available on [Pubmed] (https://pubmed.ncbi.nlm.nih.gov/31924477/).

### 2 Loading required packages

We load the packages required to perform all the analytic steps presented in this document.

```
library("DESeq2")
library("topGO")
library("pcaExplorer")
library("ideal")
library("GeneTonic")
library("apeglm")
library("dplyr")
library("msigdbr")
library("visNetwork")
library("org.Mm.eg.db")
```

### 3 Data processing

In this example we analyse data available on the Gene Expression Omnibus under
accession number GSE130842.
From the available data, we downloaded the
GSE130842\_Count\_table\_Delacher\_et\_al\_2019.xlsx Excel file, which is also
available in the Github
repository to this document.

We will preprocess the data for its use in *Gedi*
according to the workflow described in the package vignette of
*Gedi*. The package vignette is available through using
the command `browseVignette(GeDi)` after successfully installing the package.

In the first step of the preprocessing, we will generate a `DESeqDataset`
(Love, Huber, and Anders 2014) from the count table available in the Excel file. We will further
preprocess this object, called `dds_tregs`, to prepare the data for its use in
`GeDi`. We will also read in some metadata that we have set up for the dataset.
This file will also be available in the repository
as well as the final generated `dds_tregs`.

```
# We read in the data form the Excel file
count_df <- readxl::read_excel("GSE130842_Count_table_Delacher_et_al_2019.xlsx")

# We transform the data into a matrix and set the gene ids as the rownames of 
# the final matrix
count_matrix <- as.matrix(count_df[, -1])
rownames(count_matrix) <- count_df$ID

# We read in the metadata
coldata <- readxl::read_excel("GSE130842_metadata.xlsx")
coldata
```

```
## # A tibble: 56 × 5
##    ID_GEO                      group                    group2     rep_nr tissue
##    <chr>                       <chr>                    <chr>       <dbl> <chr> 
##  1 Klrg1-Il1rl1-Treg_BM_R1     Klrg1-Il1rl1-Treg_BM     Klrg1-Il1…      1 bonem…
##  2 Klrg1-Il1rl1-Treg_BM_R2     Klrg1-Il1rl1-Treg_BM     Klrg1-Il1…      2 bonem…
##  3 Klrg1-Il1rl1-Treg_BM_R3     Klrg1-Il1rl1-Treg_BM     Klrg1-Il1…      3 bonem…
##  4 Klrg1-Il1rl1-Treg_BM_R4     Klrg1-Il1rl1-Treg_BM     Klrg1-Il1…      4 bonem…
##  5 Klrg1-Il1rl1-Treg_Spleen_R1 Klrg1-Il1rl1-Treg_Spleen Klrg1-Il1…      1 spleen
##  6 Klrg1-Il1rl1-Treg_Spleen_R2 Klrg1-Il1rl1-Treg_Spleen Klrg1-Il1…      2 spleen
##  7 Klrg1-Il1rl1-Treg_Spleen_R3 Klrg1-Il1rl1-Treg_Spleen Klrg1-Il1…      3 spleen
##  8 Klrg1-Il1rl1-Treg_Spleen_R4 Klrg1-Il1rl1-Treg_Spleen Klrg1-Il1…      4 spleen
##  9 tisTregST2_BM_R1            tisTregST2_BM            tisTregST2      1 bonem…
## 10 tisTregST2_BM_R2            tisTregST2_BM            tisTregST2      2 bonem…
## # ℹ 46 more rows
```

```
# We build up the DESeqDataset object from the count data and the metadata
# As design we will choose the group of the data which indicates the tissue
# of origin as well as t5he type of Tcells in the sample
dds_tregs <- DESeqDataSetFromMatrix(
  countData = count_matrix,
  colData = coldata,
  design = ~group
)

# We transform the columnnames of the dds_tregs to include the replicate number
# of each sample
colnames(dds_tregs) <- paste0(coldata$group, "_r", coldata$rep_nr)

# Lastly we can have a look at our DESeqDataset object 
dds_tregs
```

```
## class: DESeqDataSet 
## dim: 52550 56 
## metadata(1): version
## assays(1): counts
## rownames(52550): ENSMUSG00000000001 ENSMUSG00000000003 ...
##   ENSMUSG00000114967 ENSMUSG00000114968
## rowData names(0):
## colnames(56): Klrg1-Il1rl1-Treg_BM_r1 Klrg1-Il1rl1-Treg_BM_r2 ...
##   Treg_IL2_r3 Treg_IL2_r4
## colData names(5): ID_GEO group group2 rep_nr tissue
```

Now we can see that we have set up a `DESeqDataset` object on all the available
samples. However, in this analysis, we will only focus on a subset of samples as
shown in Figure 2 of the original manuscript (Delacher et al. 2020). Hence, we will
subset our `dds_tregs` object to the groups used in the figure.

```
dds_tregs <- dds_tregs[, dds_tregs$group %in% 
                          c("KLRGminusNFIL3minusTreg" ,
                            "KLRGminusNFIL3plusTreg",
                            "KLRGplusNFIL3plusTreg",
                            "tisTregST2_BM",
                            "tisTregST2_Fat",
                            "tisTregST2_Liver",
                            "tisTregST2_Lung",
                            "tisTregST2_Skin"
                            )]

dds_tregs$group <- droplevels(dds_tregs$group)
design(dds_tregs) <- ~group
```

#### 3.1 Exploratory data analysis

In a first analysis step, we do an exploratory data analysis as described in
(Ludt et al. 2022) using the *pcaExplorer* package (Marini and Binder 2019).

We will first apply a variance-stabilizing transformation before we plot a PCA
and a sample-to-sample distance heatmap.

```
# Apply the vst transformation
vst_tregs <- vst(dds_tregs)

# Plot the PCA
pcaExplorer::pcaplot(vst_tregs,
                     intgroup = "group",
                     ntop = 1000,
                     title = "PCA plot - top 1000 most variable genes",
                     ellipse = FALSE,
                     text_labels = FALSE
                     )
```

```
# Plot a sampl-to-sample distance heatmap
pheatmap::pheatmap(as.matrix(dist(t(assay(vst_tregs)))))
```

#### 3.2 Differential expression analysis

After we have done some exploratory data analysis, we can proceed with the
differential expression analysis. We use the *DESeq2*
package (Love, Huber, and Anders 2014) for this. The False Discovery Rate is set to 5%.

Before, we run the *DESeq2* analysis, we first match the
gene ids to gene names using the *pcaExplorer* package
(Marini and Binder 2019). With this we can add the gene names to the results, which are
usually better known than gene ids. The gene names will also be later used
as “Genes” column in the input for *GeDi*.

```
# Create an annodation data frame mapping gene ids to gene names. 
anno_df <- pcaExplorer::get_annotation_orgdb(dds_tregs,"org.Mm.eg.db","ENSEMBL")
# Assign a new column SWYMBOL to the dds_tregs object, which will be later used
# as "Genes" column in the input for GeDi
rowData(dds_tregs)$SYMBOL <- anno_df$gene_name[match(rownames(dds_tregs),
                                                     anno_df$gene_id)]

# Set the false discovery rate to 5%
FDR <- 0.05

# Perform the differential gene expression analysis 
dds_tregs <- DESeq(dds_tregs)
```

After we performed the analysis, we extract the results for the condition
`KLRGplusNFIL3plusTreg vs KLRGminusNFIL3minusTreg` as we only want to compare
these two groups. Afterwards we print a summary overview of the previously
extracted results.

We also use the *ideal* package (Marini, Linke, and Binder 2020) to plot an
MA-plot of the results.

In a last step, we add the gene symbols to the resulting `DataFrame` which will
later serve as our “Genes” column in the input data to `GeDi`.

```
# Extract differentially expressed genes
# Perform contrast analysis comparing "KLRGplusNFIL3plusTreg" group to "KLRGminusNFIL3minusTreg" group
# Set a log2 fold change threshold of 1 and a significance level (alpha) of 0.05
res_tregs <- results(dds_tregs,
  contrast = c("group", "KLRGplusNFIL3plusTreg", "KLRGminusNFIL3minusTreg"),
  lfcThreshold = 1, alpha = 0.05
)

# Print a summary overview of the results
summary(res_tregs)
```

```
## 
## out of 42631 with nonzero total read count
## adjusted p-value < 0.05
## LFC > 1.00 (up)    : 683, 1.6%
## LFC < -1.00 (down) : 140, 0.33%
## outliers [1]       : 551, 1.3%
## low counts [2]     : 13054, 31%
## (mean count < 1)
## [1] see 'cooksCutoff' argument of ?results
## [2] see 'independentFiltering' argument of ?results
```

```
# Plot an MA-plot of the results
ideal::plot_ma(res_tregs, 
               ylim = c(-5, 5), 
               title = "MAplot - KLRGplusNFIL3plusTreg vs KLRGminusNFIL3minusTreg")
```

```
# Add gene symbols to the results in a column "SYMBOL"
res_tregs$SYMBOL <- rowData(dds_tregs)$SYMBOL
```

#### 3.3 Functional enrichment analysis

Following the differential expression analysis, we perform a functional enrichment
analysis using the *topGO* package (Alexa and Rahnenfuhrer 2024). Before the analysis,
we first determine the set of background genes to be used, which in our case will
be the set of expressed genes in the data. We also transform the results of our
DE analysis to fit the format expectation of the `topGOtable` function from the
*pcaExplorer* package (Marini and Binder 2019).

```
# Determine the set of background genes as all genes expressed in the dataset
geneUniverseExpr <- rowData(dds_tregs)$SYMBOL[rowSums(counts(dds_tregs)) > 0]

# Extract gene symbols from the DESeq2 results object where FDR is below 0.05
# The function deseqresult2df is used to convert the DESeq2 results to a 
# dataframe format
# FDR is set to 0.05 to filter significant results
de_symbols <- deseqresult2df(res_tregs, FDR = 0.05)$SYMBOL

# Perform Gene Ontology enrichment analysis using the topGOtable function from 
# the "pcaExplorer" package
topGO_tregs <- topGOtable(
  DEgenes = de_symbols,
  BGgenes = geneUniverseExpr,
  ontology = "BP",
  geneID = "symbol",
  addGeneToTerms = TRUE,
  mapping = "org.Mm.eg.db",
  topTablerows = 500
)
```

#### 3.4 Preparing the data for GeDi

After we have performed the functional enrichment analysis, the data is almost
ready to be used with *GeDi*. However, in its current
state the data is not yet in the correct format expected by *GeDi*.
*GeDi* expects the data to have at least two columns,
one named “Genesets” containing some form of geneset identifiers and one named
“Genes” containing a list of genes belonging to the genesets. While this is not
strictly necessary to use *GeDi* on the data, it
facilitates the use of the app as the app can be used straight away instead of
having to wait for the data to be reformated. The correct data format is however
necessary, if you want to use the data as a parameter as in `GeDi(topGO_tregs)`.

Nevertheless, we want to show you here, how to adapt the data from the
*topGO* analysis to fit the data format requirements
of *GeDi*. For this we simply have to rename the
‘GO.ID’ and the ‘genes’ column of the results as these two columns already
contain the input data in the correct format.

```
# Rename columns in the topGO_tregs dataframe
# Change the column name "GO.ID" to "Genesets"
names(topGO_tregs)[names(topGO_tregs) == "GO.ID"] <- "Genesets"

# Change the column name "genes" to "Genes"
names(topGO_tregs)[names(topGO_tregs) == "genes"] <- "Genes"
```

After we have renamed the columns, the data is now ready to be used in
*GeDi*.

### 4 Running `GeDi` on the dataset

Now we can start to explore our data using *GeDi*. For
this you can either follow the chunks in this document to prepare the data or
we can load the prepared object provided with the
repository of this
document.

```
tregs_example <- readRDS("usecase_tregs_example.RDS")
```

Once we have loaded the data, we can start the app and interactively explore
the data set using:

```
GeDi(genesets = tregs_example)
```

Once the app is started, you can have interactive guidance of the user
interface and its features by usind the introductory tours of each panel of
the app.

#### 4.1 Using `GeDi`’s functions in analysis reports

The functionality of *GeDi* can be used also as
standalone functions, to be called for example in existing analysis reports
in RMarkdown, or R scripts.

In the following chunks, we show how it is possible to call some of the functions
on the dataset presented in this document.

```
# First we extract a representation of all genes in the data
genes <- GeDi::prepareGenesetData(tregs_example)

# Then, we filter out large and generic genesets
# For this we first plot a histogram of the size of the genesets
GeDi::gsHistogram(genes, gs_names = tregs_example$Genesets, gs_description = tregs_example$Term)
```

```
# Now we filter all genesets with a size of > 200 genes

tregs_example_filtered <- tregs_example[tregs_example$Genes < 200, ]

# Next we calculate one distance score matrix for the data
mm_score <- GeDi::getMeetMinMatrix(genesets = genes)
rownames(mm_score) <- colnames(mm_score) <- tregs_example$Genesets

# We can use several plotting functions of GeDi to plot the data
GeDi::distanceHeatmap(mm_score)
```

```
GeDi::distanceDendro(mm_score)
```

```
GeDi::buildGraph(GeDi::getAdjacencyMatrix(mm_score, 0.3))
```

```
## IGRAPH 2ea70fe UN-- 500 2744 -- 
## + attr: name (v/c), title (v/x)
## + edges from 2ea70fe (vertex names):
##  [1] GO:0006954--GO:0050729 GO:0006954--GO:0032689 GO:0006954--GO:0150079
##  [4] GO:0006954--GO:0071222 GO:0006954--GO:0070098 GO:0006954--GO:0002830
##  [7] GO:0006954--GO:0032715 GO:0006954--GO:0050728 GO:0006954--GO:0002724
## [10] GO:0006954--GO:0032703 GO:0006954--GO:2000404 GO:0006954--GO:0090026
## [13] GO:0006954--GO:0002317 GO:0006954--GO:0002246 GO:0006954--GO:0032815
## [16] GO:0006954--GO:0002643 GO:0006954--GO:2000108 GO:0006954--GO:2000551
## [19] GO:0006954--GO:0045624 GO:0006954--GO:0045625 GO:0006954--GO:0051043
## [22] GO:0006954--GO:0010820 GO:0006954--GO:0072678 GO:0006954--GO:0010759
## + ... omitted several edges
```

After we have calculated distance scores, we can cluster the data and visualize
the results in several different ways.

```
# First we are clustering the data
clustering <- GeDi::clustering(mm_score, threshold = 0.3, cluster_method = "markov")

# Next we can visualize the results of the clustering
GeDi::buildClusterGraph(clustering,
                        genes, 
                        topGO_tregs$Genesets)
```

```
## IGRAPH 1fbf8fe UN-- 380 6420 -- 
## + attr: name (v/c), title (v/x), cluster (v/c)
## + edges from 1fbf8fe (vertex names):
##  [1] GO:0070098--GO:0045581 GO:0070098--GO:0006954 GO:0070098--GO:0050731
##  [4] GO:0070098--GO:0045669 GO:0070098--GO:0043433 GO:0070098--GO:0043374
##  [7] GO:0070098--GO:0070269 GO:0070098--GO:0001818 GO:0070098--GO:0072672
## [10] GO:0070098--GO:1902624 GO:0070098--GO:2000404 GO:0070098--GO:0002246
## [13] GO:0070098--GO:0002710 GO:0070098--GO:0009636 GO:0070098--GO:2000551
## [16] GO:0070098--GO:0007520 GO:0070098--GO:0071345 GO:0070098--GO:0072677
## [19] GO:0070098--GO:0036035 GO:0070098--GO:0045671 GO:0070098--GO:0045766
## [22] GO:0070098--GO:1902106 GO:0070098--GO:0016485 GO:0070098--GO:0045807
## + ... omitted several edges
```

```
GeDi::getBipartiteGraph(clustering, topGO_tregs$Genesets, genes = genes)
```

```
## IGRAPH c8ce9ff DN-B 423 380 -- 
## + attr: type (v/l), name (v/c), nodeType (v/c), shape (v/c), color
## | (v/c), title (v/c), color (e/c)
## + edges from c8ce9ff (vertex names):
##  [1] Cluster 1->GO:0150079 Cluster 1->GO:0051492 Cluster 2->GO:0097028
##  [4] Cluster 2->GO:0051384 Cluster 2->GO:0140507 Cluster 3->GO:0071347
##  [7] Cluster 3->GO:0070555 Cluster 3->GO:0007160 Cluster 4->GO:0043031
## [10] Cluster 4->GO:0045624 Cluster 4->GO:0150105 Cluster 4->GO:0042119
## [13] Cluster 4->GO:0002825 Cluster 4->GO:0034115 Cluster 4->GO:0050777
## [16] Cluster 4->GO:0045861 Cluster 4->GO:0006921 Cluster 4->GO:0030206
## [19] Cluster 4->GO:0002861 Cluster 5->GO:0050728 Cluster 5->GO:0007417
## + ... omitted several edges
```

```
GeDi::enrichmentWordcloud(topGO_tregs)
```

### Session information

```
sessionInfo()
```

```
## R version 4.5.0 (2025-04-11)
## Platform: x86_64-apple-darwin20
## Running under: macOS Sonoma 14.7
## 
## Matrix products: default
## BLAS:   /System/Library/Frameworks/Accelerate.framework/Versions/A/Frameworks/vecLib.framework/Versions/A/libBLAS.dylib 
## LAPACK: /Library/Frameworks/R.framework/Versions/4.5-x86_64/Resources/lib/libRlapack.dylib;  LAPACK version 3.12.1
## 
## locale:
## [1] en_US.UTF-8/en_US.UTF-8/en_US.UTF-8/C/en_US.UTF-8/en_US.UTF-8
## 
## time zone: Europe/Berlin
## tzcode source: internal
## 
## attached base packages:
## [1] stats4    stats     graphics  grDevices utils     datasets  methods  
## [8] base     
## 
## other attached packages:
##  [1] org.Mm.eg.db_3.21.0         visNetwork_2.1.2           
##  [3] org.Hs.eg.db_3.21.0         macrophage_1.25.0          
##  [5] msigdbr_25.1.0              dplyr_1.1.4                
##  [7] apeglm_1.31.0               GeneTonic_3.3.2            
##  [9] ideal_2.3.0                 pcaExplorer_3.3.0          
## [11] bigmemory_4.6.4             topGO_2.61.1               
## [13] SparseM_1.84-2              GO.db_3.21.0               
## [15] AnnotationDbi_1.71.0        graph_1.87.0               
## [17] DESeq2_1.49.2               SummarizedExperiment_1.39.1
## [19] Biobase_2.69.0              MatrixGenerics_1.21.0      
## [21] matrixStats_1.5.0           GenomicRanges_1.61.1       
## [23] Seqinfo_0.99.1              IRanges_2.43.0             
## [25] S4Vectors_0.47.0            BiocGenerics_0.55.0        
## [27] generics_0.1.4              knitr_1.50                 
## 
## loaded via a namespace (and not attached):
##   [1] R.methodsS3_1.8.2        dichromat_2.0-0.1        GSEABase_1.71.0         
##   [4] progress_1.2.3           wordcloud2_0.2.1         DT_0.33                 
##   [7] Biostrings_2.77.2        vctrs_0.6.5              ggtangle_0.0.7          
##  [10] digest_0.6.37            png_0.1-8                shape_1.4.6.1           
##  [13] shinyBS_0.61.1           registry_0.5-1           ggrepel_0.9.6           
##  [16] magick_2.8.7             MASS_7.3-65              reshape2_1.4.4          
##  [19] httpuv_1.6.16            foreach_1.5.2            qvalue_2.41.0           
##  [22] withr_3.0.2              xfun_0.52                ggfun_0.1.9             
##  [25] survival_3.8-3           memoise_2.0.1            clusterProfiler_4.17.0  
##  [28] gson_0.1.0               BiasedUrn_2.0.12         tidytree_0.4.6          
##  [31] GlobalOptions_0.1.2      gtools_3.9.5             R.oo_1.27.1             
##  [34] prettyunits_1.2.0        KEGGREST_1.49.1          promises_1.3.3          
##  [37] httr_1.4.7               restfulr_0.0.16          hash_2.2.6.3            
##  [40] rstudioapi_0.17.1        shinyAce_0.4.4           UCSC.utils_1.5.0        
##  [43] miniUI_0.1.2             DOSE_4.3.0               base64enc_0.1-3         
##  [46] babelgene_22.9           curl_6.4.0               polyclip_1.10-7         
##  [49] ca_0.71.1                SparseArray_1.9.0        GeDi_1.5.0              
##  [52] RBGL_1.85.0              threejs_0.3.4            xtable_1.8-4            
##  [55] stringr_1.5.1            doParallel_1.0.17        evaluate_1.0.4          
##  [58] S4Arrays_1.9.1           BiocFileCache_2.99.5     hms_1.1.3               
##  [61] bookdown_0.43            colorspace_2.1-1         filelock_1.0.3          
##  [64] NLP_0.3-2                readxl_1.4.5             shinyWidgets_0.9.0      
##  [67] magrittr_2.0.3           Rgraphviz_2.53.0         later_1.4.2             
##  [70] viridis_0.6.5            ggtree_3.17.0            lattice_0.22-7          
##  [73] NMF_0.28                 genefilter_1.91.0        XML_3.99-0.18           
##  [76] cowplot_1.2.0            pillar_1.11.0            nlme_3.1-168            
##  [79] iterators_1.0.14         gridBase_0.4-7           caTools_1.18.3          
##  [82] compiler_4.5.0           stringi_1.8.7            shinycssloaders_1.1.0   
##  [85] Category_2.75.0          TSP_1.2-5                dendextend_1.19.0       
##  [88] GenomicAlignments_1.45.1 plyr_1.8.9               crayon_1.5.3            
##  [91] abind_1.4-8              BiocIO_1.19.0            ggdendro_0.2.0          
##  [94] gridGraphics_0.5-1       emdbook_1.3.13           chron_2.3-62            
##  [97] locfit_1.5-9.12          bit_4.6.0                UpSetR_1.4.0            
## [100] fastmatch_1.1-6          codetools_0.2-20         crosstalk_1.2.1         
## [103] bslib_0.9.0              slam_0.1-55              GetoptLong_1.0.5        
## [106] tm_0.7-16                plotly_4.11.0            mime_0.13               
## [109] mosdef_1.5.1             splines_4.5.0            circlize_0.4.16         
## [112] Rcpp_1.1.0               dbplyr_2.5.0             tippy_0.1.0             
## [115] cellranger_1.1.0         utf8_1.2.6               blob_1.2.4              
## [118] clue_0.3-66              fs_1.6.6                 backbone_2.1.4          
## [121] IHW_1.37.0               expm_1.0-0               ggplotify_0.1.2         
## [124] sqldf_0.4-11             tibble_3.3.0             Matrix_1.7-3            
## [127] statmod_1.5.0            tweenr_2.0.3             pkgconfig_2.0.3         
## [130] pheatmap_1.0.13          tools_4.5.0              cachem_1.1.0            
## [133] RSQLite_2.4.1            numDeriv_2016.8-1.1      viridisLite_0.4.2       
## [136] DBI_1.2.3                fastmap_1.2.0            rmarkdown_2.29          
## [139] scales_1.4.0             grid_4.5.0               shinydashboard_0.7.3    
## [142] Rsamtools_2.25.1         sass_0.4.10              patchwork_1.3.1         
## [145] coda_0.19-4.1            BiocManager_1.30.26      fontawesome_0.5.3       
## [148] farver_2.1.2             mgcv_1.9-3               gsubfn_0.7              
## [151] yaml_2.3.10              AnnotationForge_1.51.0   rtracklayer_1.69.1      
## [154] cli_3.6.5                purrr_1.0.4              txdbmaker_1.5.6         
## [157] webshot_0.5.5            lifecycle_1.0.4          rsconnect_1.5.0         
## [160] mvtnorm_1.3-3            rintrojs_0.3.4           BiocParallel_1.43.4     
## [163] annotate_1.87.0          gtable_0.3.6             rjson_0.2.23            
## [166] ggridges_0.5.6           parallel_4.5.0           ape_5.8-1               
## [169] limma_3.65.1             jsonlite_2.0.0           colourpicker_1.3.0      
## [172] seriation_1.5.7          bitops_1.0-9             ggplot2_3.5.2           
## [175] bigmemory.sri_0.1.8      bit64_4.6.0-1            assertthat_0.2.1        
## [178] yulab.utils_0.2.0        BiocNeighbors_2.3.1      proto_1.0.0             
## [181] heatmaply_1.5.0          geneLenDataBase_1.45.0   bs4Dash_2.3.4           
## [184] bdsmatrix_1.3-7          jquerylib_0.1.4          GOSemSim_2.35.0         
## [187] R.utils_2.13.0           lazyeval_0.2.2           shiny_1.11.1            
## [190] dynamicTreeCut_1.63-1    htmltools_0.5.8.1        enrichplot_1.29.1       
## [193] rappdirs_0.3.3           STRINGdb_2.21.0          glue_1.8.0              
## [196] httr2_1.1.2              XVector_0.49.0           RCurl_1.98-1.17         
## [199] treeio_1.33.0            ComplexUpset_1.3.3       gridExtra_2.3           
## [202] igraph_2.1.4             R6_2.6.1                 tidyr_1.3.1             
## [205] gplots_3.2.0             fdrtool_1.2.18           labeling_0.4.3          
## [208] GenomicFeatures_1.61.4   cluster_2.1.8.1          rngtools_1.5.2          
## [211] bbmle_1.0.25.1           aplot_0.2.8              GenomeInfoDb_1.45.7     
## [214] plotrix_3.8-4            DelayedArray_0.35.2      tidyselect_1.2.1        
## [217] GOstats_2.75.0           ggforce_0.5.0            xml2_1.3.8              
## [220] KernSmooth_2.23-26       goseq_1.61.0             BiocStyle_2.37.0        
## [223] data.table_1.17.6        htmlwidgets_1.6.4        fgsea_1.35.4            
## [226] ComplexHeatmap_2.25.2    RColorBrewer_1.1-3       biomaRt_2.65.0          
## [229] rlang_1.1.6              uuid_1.2-1               rentrez_1.2.4           
## [232] lpsymphony_1.37.0        Cairo_1.6-2
```

### Bibliography

Alexa, Adrian, and Jorg Rahnenfuhrer. 2024. *topGO: Enrichment Analysis for Gene Ontology*.

Delacher, Michael, Charles D. Imbusch, Agnes Hotz-Wagenblatt, Jan Philipp Mallm, Katharina Bauer, Malte Simon, Dania Riegel, et al. 2020. “Precursors for Nonlymphoid-Tissue Treg Cells Reside in Secondary Lymphoid Organs and Are Programmed by the Transcription Factor BATF.” *Immunity* 52 (2): 295–312.e11. https://doi.org/10.1016/j.immuni.2019.12.002.

Love, Michael I., Wolfgang Huber, and Simon Anders. 2014. “Moderated Estimation of Fold Change and Dispersion for RNA-Seq Data with DESeq2.” *Genome Biology* 15: 550. https://doi.org/10.1186/s13059-014-0550-8.

Ludt, Annekathrin, Arsenij Ustjanzew, Harald Binder, Konstantin Strauch, and Federico Marini. 2022. “Interactive and Reproducible Workflows for Exploring and Modeling RNA-seq Data with pcaExplorer, Ideal, and GeneTonic.” *Current Protocols* 2 (4): 1–55. https://doi.org/10.1002/cpz1.411.

Marini, Federico, and Harald Binder. 2019. “pcaExplorer: an R/Bioconductor package for interacting with RNA-seq principal components.” *BMC Bioinformatics* 20 (1): 331. https://doi.org/10.1186/s12859-019-2879-1.

Marini, Federico, Jan Linke, and Harald Binder. 2020. “ideal: an R/Bioconductor package for interactive differential expression analysis.” *BMC Bioinformatics* 21 (1): 565. https://doi.org/10.1186/s12859-020-03819-5.
