## Additional File 1 and 2 for "GeDi: Simplifying Gene Set Distances for Enhanced Omics Interpretation in R/Bioconductor": GeDi_Additional_File_2.html

Using GeDi on the macrophage dataset (ERP020977)


Code 

- Show All Code
- Hide All Code
- Download Rmd

### Using GeDi on the macrophage dataset (ERP020977)

###### Annekathrin Silvia Nedwed

Institute of Medical Biostatistics, Epidemiology and Informatics (IMBEI), Mainz   
  
###### Arsenij Ustjanzew

Institute of Medical Biostatistics, Epidemiology and Informatics (IMBEI), Mainz   
  

###### Sara Salome Helbich

Institute of Immunology, University Medical Center Mainz, Mainz, Germany   
Research Center for Immunotherapy (FZI), Mainz, Germany   
  

###### Michael Delacher

Institute of Immunology, University Medical Center Mainz, Mainz, Germany   
Research Center for Immunotherapy (FZI), Mainz, Germany   
  

###### Konstantin Strauch

Institute of Medical Biostatistics, Epidemiology and Informatics (IMBEI), Mainz   
  

###### Federico Marini

Institute of Medical Biostatistics, Epidemiology and Informatics (IMBEI), Mainz   
Research Center for Immunotherapy (FZI), Mainz, Germany   
Center for Thrombosis and Hemostasis (CTH), Mainz   
  
###### 16 July 2025

### 1 About the data

The data illustrated in this document is an RNA-seq dataset, available at the European Nucleotide Archive under the accession code ERP020977 (https://www.ebi.ac.uk/ena/browser/view/ERP020977).

The data is included in the `macrophage` package available on Bioconductor (https://www.bioconductor.org/packages/release/data/experiment/html/macrophage.html). The data was generated as part of the work to identify shared quantitative trait loci (QTLs) for chromatin accessibility and gene expression in human macrophages (Alasoo et al. 2018) - the manuscript is available at https://www.nature.com/articles/s41588-018-0046-7.

### 2 Loading required packages

We load the packages required to perform all the analytic steps presented in this document.

```
library("DESeq2")
library("topGO")
library("pcaExplorer")
library("ideal")
library("GeneTonic")
library("apeglm")
library("dplyr")
library("msigdbr")
library("macrophage")
library("org.Hs.eg.db")
library("visNetwork")
```

### 3 Data processing

Before we can use the data of the `macrophage` package in `GeDi`, we first have to preprocess and analyze the data.

For this, we first obtain the data from the package and generate a `DESeqDataset`. This step is also demonstrated and described in more detail in the package vignette of `GeDi`. The vignette is accessible through `browseVignette(GeDi)` after successfully installing the package. Similar to the the vignette, we will only use and compare the Interferon gamma (INFg) treated and the naive samples. This comparison and the resulting prepared data for `Gedi` also mirror the example data available in `GeDi`.

```
# Obtain the data from the macrophage package
data("gse", package = "macrophage")

# Create a DESeqDataset object from the data choosing the line and condition
# of the samples as design.
dds_macrophage <- DESeqDataSet(gse, design = ~line + condition)

# Transform the Gencode identifiers to Ensembl identifiers
rownames(dds_macrophage) <- substr(rownames(dds_macrophage), 1, 15)

# Have a look at the object
dds_macrophage
```

```
## class: DESeqDataSet 
## dim: 58294 24 
## metadata(7): tximetaInfo quantInfo ... txdbInfo version
## assays(3): counts abundance avgTxLength
## rownames(58294): ENSG00000000003 ENSG00000000005 ... ENSG00000285993
##   ENSG00000285994
## rowData names(2): gene_id SYMBOL
## colnames(24): SAMEA103885102 SAMEA103885347 ... SAMEA103885308
##   SAMEA103884949
## colData names(15): names sample_id ... condition line
```

#### 3.1 Differential expression analysis

After we have set up the the `DESeq2` object of the `macrophage` dataset, we can follow the `DESeq2` workflow and determine the differentially expressed genes.

We use a False Discovery Rate of 5% similar to the vignette of `GeDi`.

However, before we perform the DE analysis, we filter for low expressed genes. In this example we filter all genes that do not have at least 10 counts in at least 6 samples (where 6 is the size of the smallest group in the data).

```
# Filter genes based on read counts
# Calculate the number of genes with at least 10 counts in at least 6 samples
keep <- rowSums(counts(dds_macrophage) >= 10) >= 6

# Subset the DESeqDataSet object to keep only the selected genes
dds_macrophage <- dds_macrophage[keep, ]

# Have a look at the resulting DESeqDataSet object
dds_macrophage
```

```
## class: DESeqDataSet 
## dim: 17806 24 
## metadata(7): tximetaInfo quantInfo ... txdbInfo version
## assays(3): counts abundance avgTxLength
## rownames(17806): ENSG00000000003 ENSG00000000419 ... ENSG00000285982
##   ENSG00000285994
## rowData names(2): gene_id SYMBOL
## colnames(24): SAMEA103885102 SAMEA103885347 ... SAMEA103885308
##   SAMEA103884949
## colData names(15): names sample_id ... condition line
```

```
# Set the false discovery rate to 5%
FDR <- 0.05

# Perform the differential gene expression analysis 
dds_macrophage <- DESeq(dds_macrophage)
```

After we performed the analysis, we extract the results for the condition IFNg vs naive as we only want to compare the Interferon gamma treated and the naive samples. Afterwards we print a summary overview of the previously extracted results.

In a last step, we add the gene symbols to the resulting `DataFrame` which will later serve as our Genes column in the input data to `GeDi`.

```
# Extract differentially expressed genes
# Perform contrast analysis comparing "IFNg" condition to "naive" condition
# Set a log2 fold change threshold of 1 and a significance level (alpha) of 0.05
res_macrophage_IFNg_vs_naive <- results(dds_macrophage,
  contrast = c("condition", "IFNg", "naive"),
  lfcThreshold = 1, alpha = 0.05
)

# Print a summary overview of the results
summary(res_macrophage_IFNg_vs_naive)
```

```
## 
## out of 17806 with nonzero total read count
## adjusted p-value < 0.05
## LFC > 1.00 (up)    : 652, 3.7%
## LFC < -1.00 (down) : 372, 2.1%
## outliers [1]       : 0, 0%
## low counts [2]     : 0, 0%
## (mean count < 3)
## [1] see 'cooksCutoff' argument of ?results
## [2] see 'independentFiltering' argument of ?results
```

```
# Add gene symbols to the results in a column "SYMBOL"
res_macrophage_IFNg_vs_naive$SYMBOL <- rowData(dds_macrophage)$SYMBOL
```

#### 3.2 Functional enrichment analysis

Following the differential expression analysis, we perform a functional enrichment analysis using the `topGO` package. Before the analysis, we first determine the set of background genes to be used, which in our case will be the set of expressed genes in the data. We also transform the results of our DE analysis to fit the format expectation of the `topGOtable` function.

```
# Determine the set of background genes as all genes expressed in the dataset
geneUniverseExpr <- rowData(dds_macrophage)$SYMBOL[rowSums(counts(dds_macrophage)) > 0]

# Extract gene symbols from the DESeq2 results object where FDR is below 0.05
# The function deseqresult2df is used to convert the DESeq2 results to a 
# dataframe format
# FDR is set to 0.05 to filter significant results
de_symbols_IFNg_vs_naive <- deseqresult2df(res_macrophage_IFNg_vs_naive, FDR = 0.05)$SYMBOL

# Perform Gene Ontology enrichment analysis using the topGOtable function from 
# the "pcaExplorer" package
topGO_IFNg_vs_naive <- topGOtable(
  DEgenes = de_symbols_IFNg_vs_naive,
  BGgenes = geneUniverseExpr,
  ontology = "BP",
  geneID = "symbol",
  addGeneToTerms = TRUE,
  mapping = "org.Hs.eg.db",
  topTablerows = 500
)
```

#### 3.3 Preparing the data for GeDi

After we have performed the functional enrichment analysis, the data is almost ready to be used with `GeDi`. However, in its current state the data is not yet in the correct format expected by `GeDi`. `GeDi` expects the data to have at least two columns, one named Genesets containing some form of geneset identifiers and one named Genes containing a list of genes belonging to the genesets. While this is not strictly necessary to use `GeDi` on the data, it facilitates the use of the app as the app can be used straight away instead of having to wait for the data to be reformated. The correct data format is however necessary, if you want to use the data as a parameter as in `GeDi(topGO_IFNg_vs_naive)`.

Nevertheless, we want to show you here, how to adapt the data from the `topGO` analysis to fit the data format requirements of `GeDi`. For this we simply have to rename the ‘GO.ID’ and the ‘genes’ column of the results as these two columns already contain the input data in the correct format.

```
# Rename columns in the topGO_IFNg_vs_naive dataframe
# Change the column name "GO.ID" to "Genesets"
names(topGO_IFNg_vs_naive)[names(topGO_IFNg_vs_naive) == "GO.ID"] <- "Genesets"

# Change the column name "genes" to "Genes"
names(topGO_IFNg_vs_naive)[names(topGO_IFNg_vs_naive) == "genes"] <- "Genes"
```

After we have renamed the columns, the data is now ready to be used in `GeDi`.

### 4 Running `GeDi` on the dataset

Now we can start to explore our data using `GeDi`. For this you can either follow the chunks in this document to prepare the data or we can load the prepared object provided with the repository of this document (https://github.com/AnnekathrinSilvia/GeDi\_supplement).

```
macrophage_example <- readRDS("usecase_macrophage_example.RDS")
```

Once we have loaded the data, we can start the app and interactively explore the data set using:

```
GeDi(genesets = macrophage_example)
```

Once the app is started, you can have interactive guidance of the user interface and its features by using the introductory tours of each panel of the app.

#### 4.1 Using `GeDi`’s functions in analysis reports

The functionality of `GeDi` can be used also as standalone functions, to be called for example in existing analysis reports in RMarkdown, or R scripts.

First, we will calculate the distance scores for our data. For this, we will use the Jaccard-Score to demonstrate how distance scores can be calculated. Before these score can be calculated, we have to prepare the data using the `prepareGenesetData()`function - something which is usually done internally in the app, when using `GeDi`.

```
genes <- GeDi::prepareGenesetData(macrophage_example)

jaccard_score <- GeDi::getJaccardMatrix(genesets = genes)

rownames(jaccard_score) <- colnames(jaccard_score) <- macrophage_example$Genesets
```

Afterwards, we can use the distance scores to generate some plots of the calculated scores, such as a heatmap or a network.

```
GeDi::distanceHeatmap(jaccard_score)
```

```
graph <- GeDi::buildGraph(GeDi::getAdjacencyMatrix(jaccard_score, cutOff = 0.3))
  
visNetwork::visIgraph(graph) %>%
          visNodes(color = list(
            background = "#0092AC",
            highlight = "gold",
            hover = "gold"
          )) %>%
          visEdges(color = list(
            background = "#0092AC",
            highlight = "gold",
            hover = "gold"
          )) %>%
          visOptions(
            highlightNearest = list(
              enabled = TRUE,
              degree = 1,
              hover = TRUE
            ),
            nodesIdSelection = TRUE
          ) %>%
          visExport(
            name = "distance_scores_network",
            type = "png",
            label = "Save Distance Scores graph"
          )
```

Once distance scores are calculated, `GeDi`can be used to cluster the genesets based on their calculated pairwise similarity scores. In this example, we will use the Louvain clustering algorithm for this. Afterwards, we use `GeDi`’s visualisation options to visualise the clustering results.

```
louvain_clustering <- GeDi::louvainClustering(jaccard_score, threshold = 0.3)
```

```
graph <- GeDi::buildClusterGraph(louvain_clustering, 
                                 macrophage_example,
                                 macrophage_example$Genesets, 
                                 color_by = "Cluster",
                                 gs_names = macrophage_example$Term)

visNetwork::visIgraph(graph) %>%
          visOptions(
            highlightNearest = list(
              enabled = TRUE,
              degree = 1,
              hover = TRUE
            ),
            nodesIdSelection = TRUE,
            selectedBy = list(variable = "cluster", multiple = TRUE)
          ) %>%
          visExport(
            name = "cluster_network",
            type = "png",
            label = "Save Cluster graph"
          )
```

```
# Plot an enrichment wordcloud for the first cluster
GeDi::enrichmentWordcloud(macrophage_example[louvain_clustering[[1]],])
```

### Session information

```
sessionInfo()
```

```
## R version 4.5.0 (2025-04-11)
## Platform: x86_64-apple-darwin20
## Running under: macOS Sonoma 14.7
## 
## Matrix products: default
## BLAS:   /System/Library/Frameworks/Accelerate.framework/Versions/A/Frameworks/vecLib.framework/Versions/A/libBLAS.dylib 
## LAPACK: /Library/Frameworks/R.framework/Versions/4.5-x86_64/Resources/lib/libRlapack.dylib;  LAPACK version 3.12.1
## 
## locale:
## [1] en_US.UTF-8/en_US.UTF-8/en_US.UTF-8/C/en_US.UTF-8/en_US.UTF-8
## 
## time zone: Europe/Berlin
## tzcode source: internal
## 
## attached base packages:
## [1] stats4    stats     graphics  grDevices utils     datasets  methods  
## [8] base     
## 
## other attached packages:
##  [1] visNetwork_2.1.2            org.Hs.eg.db_3.21.0        
##  [3] macrophage_1.25.0           msigdbr_25.1.0             
##  [5] dplyr_1.1.4                 apeglm_1.31.0              
##  [7] GeneTonic_3.3.2             ideal_2.3.0                
##  [9] pcaExplorer_3.3.0           bigmemory_4.6.4            
## [11] topGO_2.61.1                SparseM_1.84-2             
## [13] GO.db_3.21.0                AnnotationDbi_1.71.0       
## [15] graph_1.87.0                DESeq2_1.49.2              
## [17] SummarizedExperiment_1.39.1 Biobase_2.69.0             
## [19] MatrixGenerics_1.21.0       matrixStats_1.5.0          
## [21] GenomicRanges_1.61.1        Seqinfo_0.99.1             
## [23] IRanges_2.43.0              S4Vectors_0.47.0           
## [25] BiocGenerics_0.55.0         generics_0.1.4             
## [27] knitr_1.50                 
## 
## loaded via a namespace (and not attached):
##   [1] R.methodsS3_1.8.2        dichromat_2.0-0.1        GSEABase_1.71.0         
##   [4] progress_1.2.3           wordcloud2_0.2.1         DT_0.33                 
##   [7] Biostrings_2.77.2        vctrs_0.6.5              ggtangle_0.0.7          
##  [10] digest_0.6.37            png_0.1-8                shape_1.4.6.1           
##  [13] shinyBS_0.61.1           registry_0.5-1           ggrepel_0.9.6           
##  [16] magick_2.8.7             MASS_7.3-65              reshape2_1.4.4          
##  [19] httpuv_1.6.16            foreach_1.5.2            qvalue_2.41.0           
##  [22] withr_3.0.2              xfun_0.52                ggfun_0.1.9             
##  [25] survival_3.8-3           memoise_2.0.1            clusterProfiler_4.17.0  
##  [28] gson_0.1.0               BiasedUrn_2.0.12         tidytree_0.4.6          
##  [31] GlobalOptions_0.1.2      gtools_3.9.5             R.oo_1.27.1             
##  [34] prettyunits_1.2.0        KEGGREST_1.49.1          promises_1.3.3          
##  [37] httr_1.4.7               restfulr_0.0.16          hash_2.2.6.3            
##  [40] rstudioapi_0.17.1        shinyAce_0.4.4           UCSC.utils_1.5.0        
##  [43] miniUI_0.1.2             DOSE_4.3.0               base64enc_0.1-3         
##  [46] babelgene_22.9           curl_6.4.0               polyclip_1.10-7         
##  [49] ca_0.71.1                SparseArray_1.9.0        GeDi_1.5.0              
##  [52] RBGL_1.85.0              threejs_0.3.4            xtable_1.8-4            
##  [55] stringr_1.5.1            doParallel_1.0.17        evaluate_1.0.4          
##  [58] S4Arrays_1.9.1           BiocFileCache_2.99.5     hms_1.1.3               
##  [61] bookdown_0.43            colorspace_2.1-1         filelock_1.0.3          
##  [64] NLP_0.3-2                shinyWidgets_0.9.0       magrittr_2.0.3          
##  [67] Rgraphviz_2.53.0         later_1.4.2              viridis_0.6.5           
##  [70] ggtree_3.17.0            lattice_0.22-7           NMF_0.28                
##  [73] genefilter_1.91.0        XML_3.99-0.18            cowplot_1.2.0           
##  [76] pillar_1.11.0            nlme_3.1-168             iterators_1.0.14        
##  [79] gridBase_0.4-7           caTools_1.18.3           compiler_4.5.0          
##  [82] stringi_1.8.7            shinycssloaders_1.1.0    Category_2.75.0         
##  [85] TSP_1.2-5                dendextend_1.19.0        GenomicAlignments_1.45.1
##  [88] plyr_1.8.9               crayon_1.5.3             abind_1.4-8             
##  [91] BiocIO_1.19.0            ggdendro_0.2.0           gridGraphics_0.5-1      
##  [94] emdbook_1.3.13           chron_2.3-62             locfit_1.5-9.12         
##  [97] bit_4.6.0                UpSetR_1.4.0             fastmatch_1.1-6         
## [100] codetools_0.2-20         crosstalk_1.2.1          bslib_0.9.0             
## [103] slam_0.1-55              GetoptLong_1.0.5         tm_0.7-16               
## [106] plotly_4.11.0            mime_0.13                mosdef_1.5.1            
## [109] splines_4.5.0            circlize_0.4.16          Rcpp_1.1.0              
## [112] dbplyr_2.5.0             tippy_0.1.0              blob_1.2.4              
## [115] clue_0.3-66              fs_1.6.6                 backbone_2.1.4          
## [118] IHW_1.37.0               expm_1.0-0               ggplotify_0.1.2         
## [121] sqldf_0.4-11             tibble_3.3.0             Matrix_1.7-3            
## [124] statmod_1.5.0            tweenr_2.0.3             pkgconfig_2.0.3         
## [127] pheatmap_1.0.13          tools_4.5.0              cachem_1.1.0            
## [130] RSQLite_2.4.1            numDeriv_2016.8-1.1      viridisLite_0.4.2       
## [133] DBI_1.2.3                fastmap_1.2.0            rmarkdown_2.29          
## [136] scales_1.4.0             grid_4.5.0               shinydashboard_0.7.3    
## [139] Rsamtools_2.25.1         sass_0.4.10              patchwork_1.3.1         
## [142] coda_0.19-4.1            BiocManager_1.30.26      fontawesome_0.5.3       
## [145] farver_2.1.2             mgcv_1.9-3               gsubfn_0.7              
## [148] yaml_2.3.10              AnnotationForge_1.51.0   rtracklayer_1.69.1      
## [151] cli_3.6.5                purrr_1.0.4              txdbmaker_1.5.6         
## [154] webshot_0.5.5            lifecycle_1.0.4          rsconnect_1.5.0         
## [157] mvtnorm_1.3-3            rintrojs_0.3.4           BiocParallel_1.43.4     
## [160] annotate_1.87.0          gtable_0.3.6             rjson_0.2.23            
## [163] ggridges_0.5.6           parallel_4.5.0           ape_5.8-1               
## [166] limma_3.65.1             jsonlite_2.0.0           colourpicker_1.3.0      
## [169] seriation_1.5.7          bitops_1.0-9             ggplot2_3.5.2           
## [172] bigmemory.sri_0.1.8      bit64_4.6.0-1            assertthat_0.2.1        
## [175] yulab.utils_0.2.0        BiocNeighbors_2.3.1      proto_1.0.0             
## [178] heatmaply_1.5.0          geneLenDataBase_1.45.0   bs4Dash_2.3.4           
## [181] bdsmatrix_1.3-7          jquerylib_0.1.4          GOSemSim_2.35.0         
## [184] R.utils_2.13.0           lazyeval_0.2.2           shiny_1.11.1            
## [187] dynamicTreeCut_1.63-1    htmltools_0.5.8.1        enrichplot_1.29.1       
## [190] rappdirs_0.3.3           STRINGdb_2.21.0          glue_1.8.0              
## [193] httr2_1.1.2              XVector_0.49.0           RCurl_1.98-1.17         
## [196] treeio_1.33.0            ComplexUpset_1.3.3       gridExtra_2.3           
## [199] igraph_2.1.4             R6_2.6.1                 tidyr_1.3.1             
## [202] gplots_3.2.0             fdrtool_1.2.18           GenomicFeatures_1.61.4  
## [205] cluster_2.1.8.1          rngtools_1.5.2           bbmle_1.0.25.1          
## [208] aplot_0.2.8              GenomeInfoDb_1.45.7      plotrix_3.8-4           
## [211] DelayedArray_0.35.2      tidyselect_1.2.1         GOstats_2.75.0          
## [214] ggforce_0.5.0            xml2_1.3.8               KernSmooth_2.23-26      
## [217] goseq_1.61.0             BiocStyle_2.37.0         data.table_1.17.6       
## [220] htmlwidgets_1.6.4        fgsea_1.35.4             ComplexHeatmap_2.25.2   
## [223] RColorBrewer_1.1-3       biomaRt_2.65.0           rlang_1.1.6             
## [226] uuid_1.2-1               rentrez_1.2.4            lpsymphony_1.37.0       
## [229] Cairo_1.6-2
```
